# Deletion or inhibition of astrocytic transglutaminase 2 promotes functional recovery after spinal cord injury

**DOI:** 10.1101/2021.09.15.460455

**Authors:** Anissa Elahi, Jacen Emerson, Jacob Rudlong, Jeffrey W. Keillor, Garrick Salois, Adam Visca, Peter Girardi, Gail V.W. Johnson, Christoph Pröschel

## Abstract

Following CNS injury astrocytes become “reactive” and exhibit pro-regenerative or harmful properties. However, the molecular mechanisms that cause astrocytes to adopt either phenotype are not well understood. Transglutaminase 2 (TG2) plays a key role in regulating the response of astrocytes to insults. Here we used mice in which TG2 was specifically deleted in astrocytes (*Gfap*-Cre+/-*TG2*fl/fl, referred to here as TG2-A-cKO) in a spinal cord contusion injury (SCI) model. Deletion of TG2 from astrocytes resulted in a significant improvement in motor function following SCI. GFAP and NG2 immunoreactivity, as well as number of SOX9 positive cells, were significantly reduced in TG2-A-cKO_mice. RNA-seq analysis of spinal cords from TG2-A-cKO and control mice 3 days postinjury identified thirty-seven differentially expressed genes, all of which were increased in TG2-A-cKO mice. Pathway analysis reveals a prevalence for fatty acid metabolism, lipid storage and energy pathways, which play essential roles in neuron-astrocyte metabolic coupling. Excitingly, treatment of wild type mice with the selective TG2 inhibitor VA4 significantly improved functional recovery after SCI, similar to what was observed using the genetic model. These findings indicate the use of TG2 inhibitors as a novel strategy for the treatment of SCI and other CNS injuries.

## 1. Introduction

Astrocytes are the most abundant cell type in the central nervous system (CNS) and play an essential role in supporting neuron homeostasis and function [1]. For example, as a key component of the tripartite synapse (which also includes presynaptic and postsynaptic nerve terminals), astrocytes play a pivotal role in modulating synaptic transmission and plasticity [1, 2]. Astrocytes also play a central role mediating the response of the CNS to injury. Following an insult, astrocytes respond by becoming reactive and undergoing a rapid change in morphology, gene expression, and molecular functions [3, 4]. Depending on the type, severity and location of the injury, astrocytes exhibit different responses, taking on a phenotype that is either beneficial or detrimental to neuron survival and axonal regeneration [5]. Although it is clear that astrocytes can take on either supportive or harmful phenotypes after injury, the cell autonomous determinants that direct astrocytes towards either phenotype have not been well defined. Recently, we have identified transglutaminase 2 (TG2) as a key factor in determining the molecular response of astrocytes to injury [6-9].

TG2 is a multifunctional protein that is upregulated in astrocytes in response to cell stressors including ischemia and inflammatory signals. Several studies have demonstrated that TG2 plays a role in facilitating either cell death or survival in a range of cell types cell types, depending on the cell type and stressor [8]. For example, TG2 supports neuronal health [10]. Overexpression of TG2 in neurons improves their survival following an ischemic insult and conversely when TG2 is depleted from neurons it significantly decreases their survival [11, 12]. In contrast, deletion or depletion of TG2 from astrocytes significantly increases their survival in response to oxygen and glucose deprivation (OGD) and significantly increases their ability to protect neurons from ischemic-induced cell death injury [6, 7, 13]. In addition, we have demonstrated that mice with astrocyte-specific TG2 deletion (glial fibrillary acidic protein (*Gfap*)-Cre+/−/TG2fl/fl, referred to here as TG2-A-cKO) exhibited significantly less astrocytic scarring in an in vivo model of a contusion spinal cord injury (SCI) when compared to TG2fl/fl mice expressing normal levels of astrocytic TG2 [7]. Given that the reduction of astrocytic scarring in the TG2-A-cKO mouse is indicative of a reduction in reactive astrogliosis in the mice with TG2-depleted astrocytes, it is important to understand how deletion of TG2 impacts functional recovery following SCI.

In this study we demonstrate that astrocytic-specific TG2 deletion results in a significant improvement in functional recovery following a T9 spinal cord contusion injury. GFAP immunoreactivity was significantly decreased in the TG2-A-cKO compared to TG2fl/fl mice. We further extend these findings and demonstrate that expression of both the transcription factor Sox9 and the chondroitin sulfate proteoglycan (CSPG) NG2 (Cspg4) are significantly decreased in TG2-A-cKO spinal cord compared to TG2fl/fl mice at 3 and 7 days after injury. These are exciting findings as NG2 is part of the astroglial scar and may block successful CNS regeneration [14], while deletion of Sox9 improves outcomes after CNS injury [15-17].

To begin to understand the molecular mechanisms contributing to improved outcomes follow SCI when TG2 is specifically deleted from astrocytes, we carried out RNA-seq on spinal cords from TG2-A-cKO and TG2fl/fl mice 3 days after SCI. Intriguingly, 37 genes were differentially expressed after injury in TG2-A-cKO spinal cords compared to TG2fl/fl mice and all were upregulated. Interestingly, the majority of genes that were upregulated were involved in fatty acid metabolism, energy pathways and lipid storage. Overall, these genes are involved in promoting neuron-astrocyte metabolic coupling which enhances the ability of the astrocytes to protect and support the neurons [18-20].

Finally, to test whether acute inhibition of TG2 activity after SCI is sufficient to promote recovery, we used a pharmacological approach. *In vitro*, the highly specific and irreversible TG2 transglutaminase inhibitor VA4 [21] phenocopies the effects of TG2 deletion in that it protects astrocytes from OGD-induced cell death [13]. In addition, previous studies with the parent compound of VA4 showed that it was well-tolerated by mice [22]. Interestingly, in mice expressing normal levels of TG2, treatment with VA4 significantly improved functional recovery following SCI compared to mice treated with vehicle only. Overall, the results of our studies demonstrate that removal of astrocytic TG2 or inhibition of TG2 significantly improves outcomes after CNS injury and strongly suggest that the use of TG2 inhibitors maybe a therapeutic strategy for treating acute SCI.

## 2. Materials and Methods

### Mice

All procedures with animals were in accordance with guidelines established by the University of Rochester Committee on Animal Resources with approval from the Institutional Animal Care and Use Committee. TG2fl/fl mice were generously provided by Drs. R. Graham and S. Iismaa (Victor Chang Cardiac Research Institute). These mice contain loxP sites flanking *Tg2* exons 6–8 [23]. B6·Cg-Tg(*Gfap* Cre)73.12Mvs/J (*Gfap* Cre+/−) mice were obtained from Jackson Laboratories and have been described and characterized previously [24]. See [7] for detailed information on the generation and genotyping of the *Gfap* Cre+/− TG2fl/fl (TG2-A-cKO) and TG2fl/fl mice.

### Mouse spinal cord contusion injury

All mice were genotyped prior to surgery and after tissue harvest. For the locomotor evaluation, female mice were used for studies that continued for 42 days. For all other measures both male and female mice were used. TG2-A-cKO and TG2fl/fl mice, 12–16 weeks of age, were anesthetized using a combination of ketamine and xylazine given intraperitoneally. A laminectomy was performed at the thoracic (T9) level and the dura mater was exposed. The spinal cord was moderately contused bilaterally using a force-defined injury device (65 kDyn, Infinite Horizon Impactor 400, Precision Systems and Instrumentation). Muscle and skin were closed in layers at the injury. A heating pad was used to maintain body temperature at 37°C during surgery and the initial recovery period. Mice were given buprenorphine subcutaneously at time of surgery and three times daily for 3 days thereafter. Food and DietGel® Recovery was placed in the bottom of the cages and lactated Ringer’s solution was administrated i.p. for the first 5 days to ensure hydration. Following SCI, bladders were manually expressed every 12 h until spontaneous voiding recurred.

For the studies with the TG2 inhibitor VA4 [21], TG2fl/fl mice expressing normal levels of TG2 were used. One hour after SCI they were administered VA4 at 15 mg/kg i.p. in Cremophor or Cremophor only (as a control) and then again 24 and 48 hours later. Mice were maintained after injury as described above.

### Locomotor evaluation

For gait analysis, mice were video recorded traversing an illuminated, 80 cm long glass walkway. A minimum of 3 complete crossings per session were recorded and two individuals blinded to the genotype of the mice viewed the videos and scored the motor function of the hind limbs using the Toyama Mouse Score (TMS) [25]. Movements of the left and right hind limbs were evaluated independently. The scores were averaged for each animal. Scoring of motor function was performed pre-injury, and then at: 1, 3, 7, 14, 21, 32 and 42 days post-injury (dpi).

### Histology

Animals were anesthetized and transcardially perfused on the indicated days post injury with 0.1 M phosphate-buffered saline (PBS) followed by 4% paraformaldehyde (PFA) in 0.1M PBS. Dissected spinal cords were cryo-protected by sequential equilibration in 10%, 20% and 30% sucrose. Transverse sections (20 μm) of spinal cords were collected rostral, central and caudal of injury site. For immunofluorescence staining, sections were washed in PBS, post fixed with 4% PFA for 10 minutes, washed 3 × 5 minutes with PBS, permeabilized with 0.5% Triton X-100 in PBS for 15 minutes, blocked with 10% goat or donkey serum, 1% bovine serum albumin, and 0.2% Triton X-100 in PBS for 1 hour at room temperature, then stained with: rabbit polyclonal (pAb) glial fibrillary acidic protein (GFAP,1:1000, Dako, Z033401-2), goat pAb glial fibrillary acidic protein (GFAP,1:300, Novus, NB100-53809), rabbit pAb SOX9 (1:50, Millipore Sigma, AB5535), chicken pAb NeuN (1:2000, Millipore Sigma, ABN91), or Anti-NG2, Alexa Fluor 488 conjugate antibody (1:100, Millipore Sigma, AB5320A4) overnight at 4°C in 0.1 M PBS, 1% goat or donkey serum, 0.2% Triton X-100 (antibody buffer). The next day sections were rinsed in PBS for 3 × 5 minutes, then sections were incubated with donkey anti-goat Alexa Fluor 488 (1:1000, Invitrogen, A-11055), goat anti-chicken Alexa Fluor 488 (1:1000, Invitrogen, A32931), goat anti-rabbit Alexa Fluor 647 (1:1000, Invitrogen, A32733), donkey anti-goat Alexa Flour 647 (1:1000, Invitrogen, A32849), donkey anti-sheep Alexa Fluor 647 (1:1000, Invitrogen, A32849) or donkey anti-rabbit Alexa Fluor 647 (1:1000, Invitrogen, A48258) as needed. Tissue sections were counterstained with 1 μg/ml DAPI (Jackson ImmunoResearch) in antibody buffer for 30 minutes. Sections were rinsed in Milli-Q water for 3 × 5 minutes and cover slipped with ProLong Gold Antifade (Invitrogen, P36930).

For immunofluorescence imaging and quantification, at least three sections per animal at 2 mm rostral (−2 mm), epicenter, and 2 mm caudal (+2 mm) of injury site were collected, stained and imaged. All sections used for signal quantification were processed simultaneously, using the same batch of reagents. Images were acquired using a TCS SP5 laser scanning confocal microscope (Leica Microsystem) with consistent settings across conditions. All analyses were performed in sequential scanning mode to prevent cross-bleeding between channels. Quantification of intensity and cell counts was performed using the Measurement and Analyze Particles functions in FIJI software (version 1.41, National Institutes of Health, USA) (supplementary code 1 & 2.ijm)(Supplemental Figure S1) [26].

### RNA isolation and cDNA synthesis

Animals were anesthetized and transcardially perfused on 3 dpi with 0.1M PBS. Spinal cords were removed and an 8 mm section (4mm rostral to caudal of lesion center) was collected and snap frozen in liquid nitrogen. Total RNA was isolated following homogenization in Trizol as per manufacturer’s instructions. Samples were then collected and spun through a HighBind® RNA spin column (Omega Biotek) and processed according to the manufacturer’s instructions (E.Z.N.A.® Total RNA Kit I). RNA was eluted in 35 μl RNAse-free water.

### RNA sequencing and analysis

RNA-seq was performed at the University of Rochester Genomics Research Center using RNA isolated from the spinal cords of 4 TG2-A-cKO and 4 TG2fl/fl mice collected 3 dpi. Analysis was also performed on 3 TG2-A-cKO and 3 TG2fl/fl mice without injury. RNA concentration was determined with the NanopDrop 1000 spectrophotometer (NanoDrop, Wilmington, DE) and RNA quality was assessed with the Agilent Bioanalyzer (Agilent, Santa Clara, CA). The TruSeq RNA Sample Preparation Kit V2 (Illumina, San Diego, CA) was used for next generation sequencing library construction according to the manufacturer’s protocols. In brief, mRNA was purified from 100 ng total RNA with oligo-dT magnetic beads and fragmented. First-strand cDNA synthesis was performed with random hexamer priming followed by second-strand cDNA synthesis. End repair and 3′ adenylation was then performed on the double stranded cDNA. Illumina adaptors were ligated to both ends of the cDNA, purified by gel electrophoresis and amplified with PCR primers specific to the adaptor sequences to generate amplicons of approximately 200–500 bp in size. The amplified libraries were hybridized to the Illumina single end flow cell and amplified using the cBot (Illumina, San Diego, CA) at a concentration of 8 pM per lane. Single end reads of 100 nt were generated for each sample. Sequenced reads were cleaned according to a rigorous pre-processing workflow (Trimmomatic-0.32) before mapping them to mouse reference genome (GRCm38.p4) with STAR-2.4.2a (https://github.com/alexdobin/STAR). Cufflinks2.0.2 (cuffdiff2 - Running Cuffdiff) was used with the gencode M6 mouse gene annotations to perform differential expression analysis with an FDR cutoff of 0.05 (95% confidence interval). Preliminary evaluation of the RNA-seq data was carried using Functional Enrichment Analysis Tool (http://www.funrich.org/download/) [27], STRING (https://string-db.org/) and gene ontology pathway analysis with Enrichr [28-30].

### Quantitative-PCR (qPCR)

First strand cDNA was reverse transcribed from 500 ng total RNA using Verso cDNA Synthesis Kit (Thermo Scientific, AB1453B). Real-time PCR was performed using PerfeCTa® SYBR® Green FastMix® PCR Reagent (Quantabio, Beverly, MA, USA) on a CFX96 Real-Time System (Bio-Rad Laboratories, Hercules, CA, USA). The primers used in this study were as follows: lipoprotein lipase (Lpl), forward 5’GGGAGTTTGGCTCCAGAGTTT3′, reverse 5’TGTGTCTTCAGGGGTCCTTAG3’; Fatty Acid Binding Protein 4 (Fabp4), forward 5’AAGGTGAAGAGCATCATAACCCT3’, reverse 5’ TCACGCCTTTCATAACACATTCC3’; Perilipin 1 (Plin1), forward 5’GGGACCTGTGAGTGCTTCC3’, reverse 5’GTATTGAAGAGCCGGGATCTTTT3’ and Glyceraldehyde-3-Phosphate Dehydrogenase (Gapdh), forward 5’ATGGGACGATGCTGGTACTAG3’, reverse 5’TGCTGACAACCTTGAGTGAAAT3’. Relative expression of target transcripts was normalized to Gapdhas a housekeeping control using ΔΔCt method.

### Statistical analysis

Data were graphed and analyzed using GraphPad Prism software. Statistical significance was tested by two-way ANOVA followed by Bonferroni’s post hoc comparisons of TG2-A-cKO vs TG2fl/fl and VA4 vs Control data sets. Data are expressed as mean ± SEM. The level of significance was set to less than 0.05.

## 3. Results

### 3.1 Deletion of TG2 from astrocytes significantly improves functional recovery after SCI

For these studies mice were scored using the TMS scale before injury and then at the indicated days after injury up to 42 days [25]. Using the TMS scale, each blinded observer was able isolate each attribute of gait, score each attribute and assign a score for functional recovery. TMS categories include frequence and mobile extent of ankle movement; movement in knee and hip; sole touch at rest and in step, coordination, parallel or rotative movement in step, and trunk stability. Each scorer totaled their score for the left hind limb and right hind limb. Both scores were averaged for the final value. The results of these measures are shown in Figure 1 and clearly demonstrate that deletion of TG2 from astrocytes significantly improves functional recovery. By day 7, TG2-A-cKO mice demonstrate a significantly higher TMS score, with an average of 5 points higher than TG2fl/fl mice. While TG2fl/fl mice also demonstrate a gradual improvement over time, animals with astrocyte-specific deletion of TG2 achieve and maintain a significantly higher TMS score through day 42 post injury.

**Figure 1:**
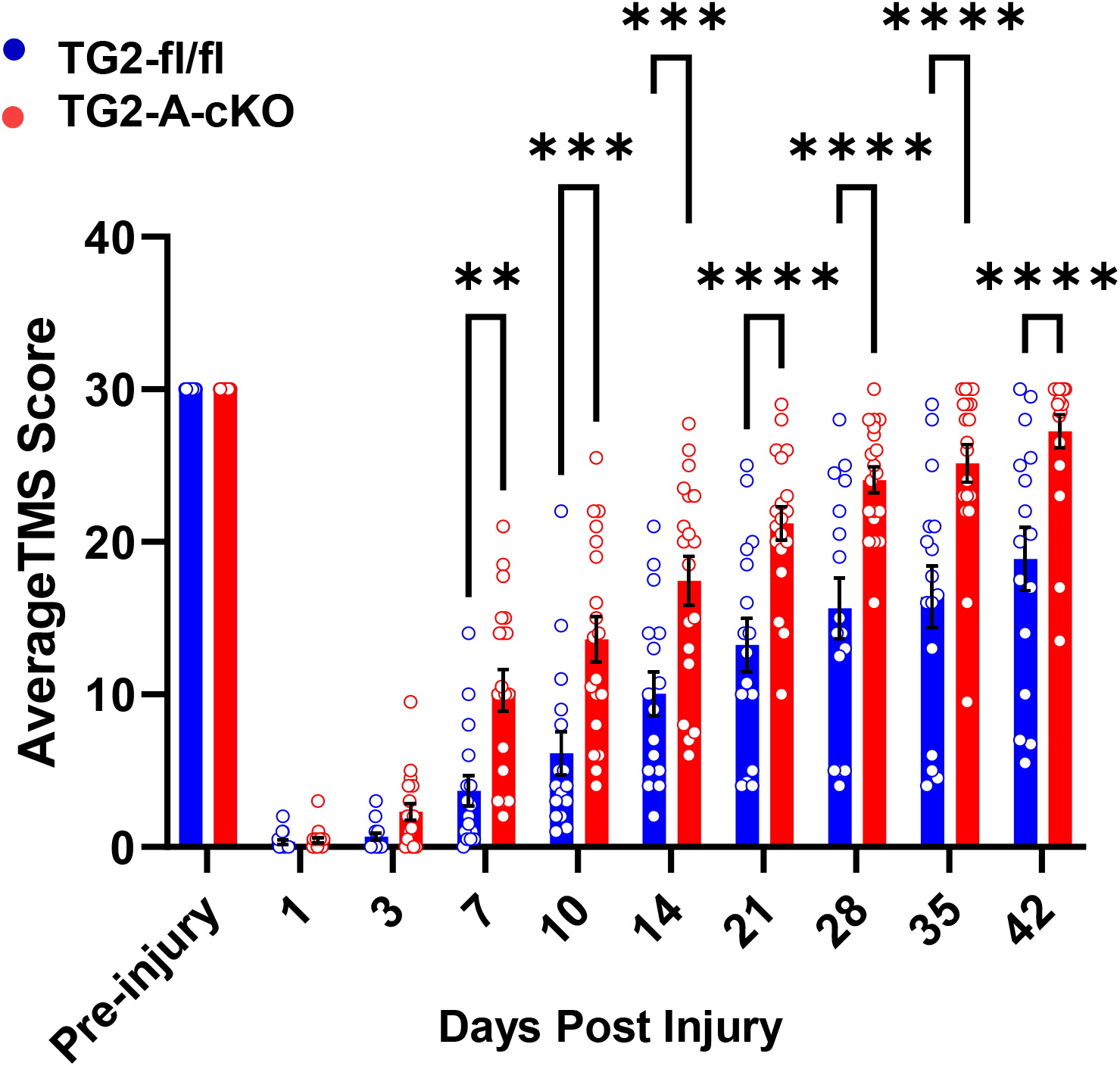
Deletion of TG2 from astrocytes improves functional recovery after spinal cord injury (SCI). TG2fl/fl and TG2-A-cKO mice at 12–16 weeks of age (12-15 mice in each group) were anesthetized and their spinal cords moderately contused blilaterally at T9. Motor function was assessed prior to injury and at the days indicated post-injury using the Toyama mouse score (TMS) [25]. Data are expressed as an average of the right hind limb and left hind limb. Data were analyzed by two-way ANOVA with a posthoc analysis using Bonferroni’s multiple comparisons test. **p<0.01, ***p<0.001, ****p<0.0001.

### 3.2 Deletion of TG2 from astrocytes results in a significant attenuation of reactive gliosis after SCI

As TG2 is rapidly upregulated in astrocytes in response to CNS injury [9], we next examined histological measures of CNS injury and reactive gliosis in spinal cord sections from mice at 3 and 7 dpi. GFAP immunoreactivity was significantly reduced both 2 mm proximal (−2 mm) and at the epicenter of the injury in TG2-A-cKO mice compared to TG2fl/fl mice that express normal levels of astrocytic TG2. This decrease in immunoreactivity was observed at both 3 and 7 dpi (**Fig. 2**). We also found that immunoreactivity of another important component of the glial scar, the chondroitin sulfate proteoglycan NG2 [14], was significantly reduced in the lesions of TG2-A-cKO mice as compared to TG2 expressing mice (**Fig. 3**). This reduction in glial scar CSPG was born out by a reduction in the number of SOX9 positive astrocytes. SOX9 is a transcription factor that upregulates genes associated with glial scar formation [16], including CSPGs such as NG2 (Cspg4) [31]. The number of SOX9 expressing cells, was reduced both at 3 and 7dpi at the epicenter of injured TG2-A-cKO mice; numbers were also significantly lower at 3 dpi at 2 mm rostral and at 7 dpi 2 mm caudal from the injury site (**Fig. 4**).

**Figure 2:**
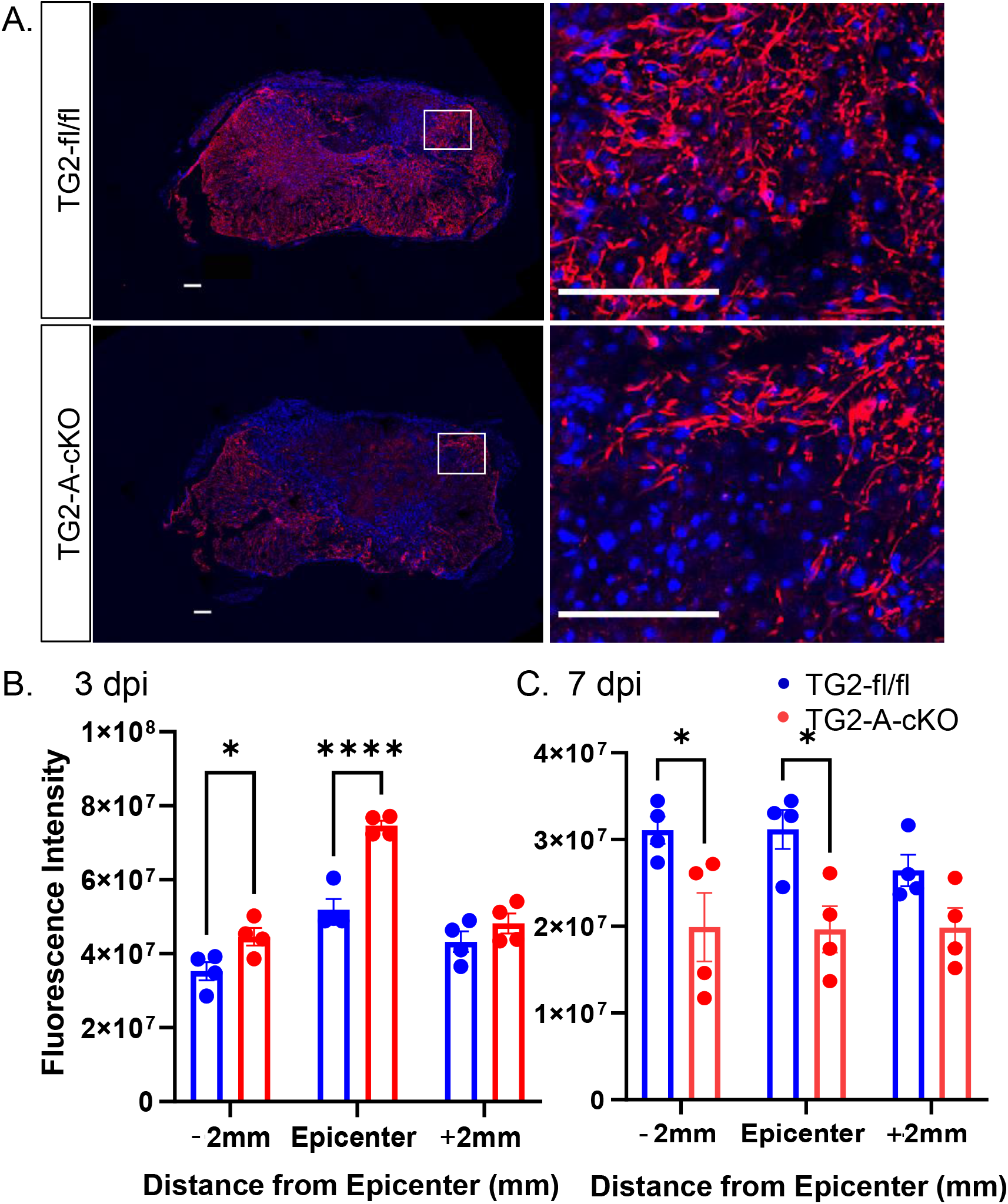
GFAP Expression is significantly reduced at the lesion site in TG2-A-cKO at 3 and 7 days after spinal cord contusion injury. **A**. Representative images of 7 days post injury (7 dpi) at the epicenter of the injury. Spinal cord sections were immunostained for GFAP (red) and counterstained with DAPI (blue). At the right are enlargements of the boxed areas in the images at the left. Scale bars = 100 μm. **B**. Quantified GFAP immunoreactivity at 3 dpi and, **C**. 7 dpi. Gfap expression was significantly lower -2 mm and at the epicenter in the TG2-A-cKO mice at both 3 and 7dpi. N=4 animals per group. *p < 0.05, ****p<0.0001.

**Figure 3:**
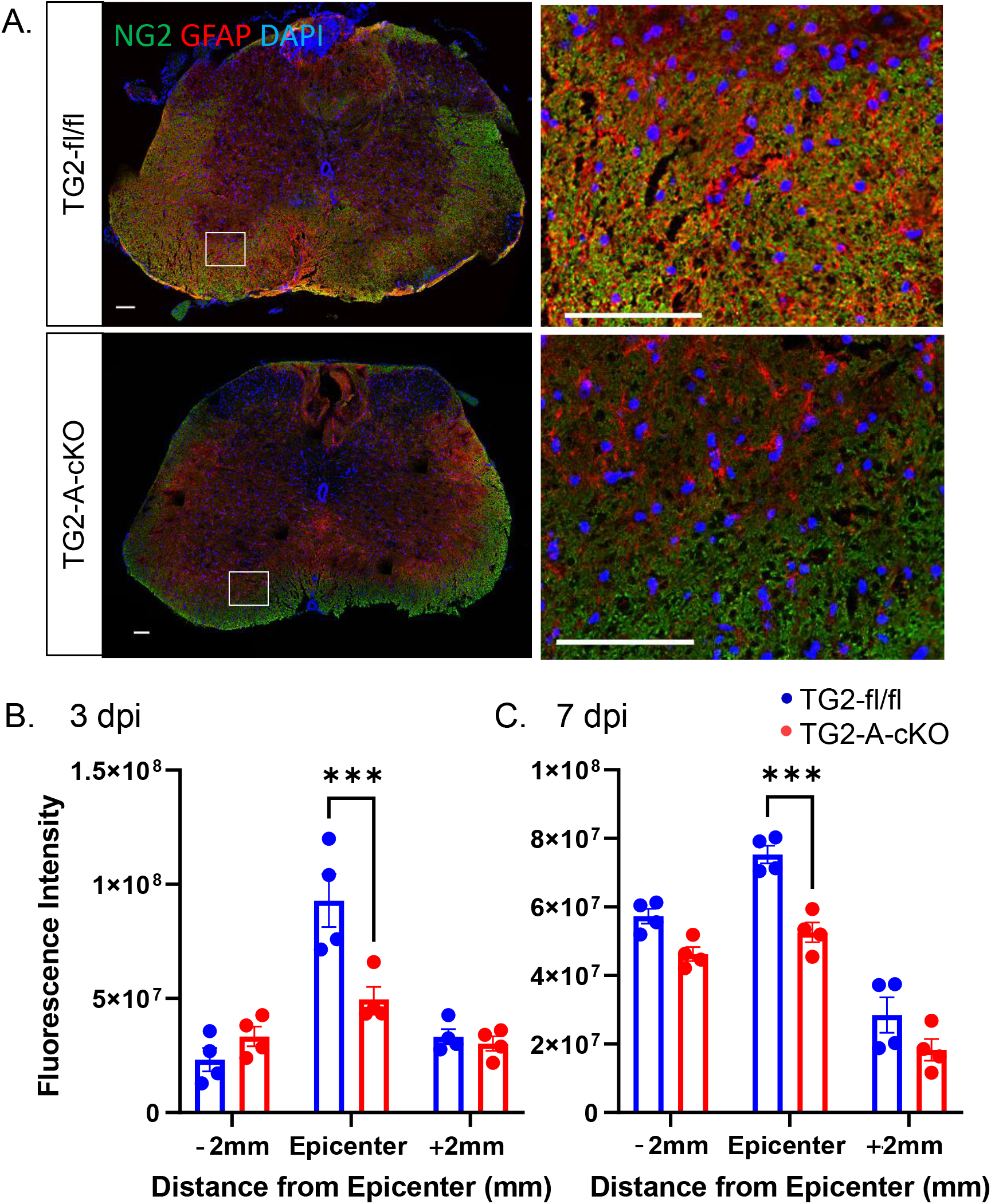
NG2 immunoreactivity is significantly reduced at the lesion site in TG2-A-cKO mice 3 and 7 days after spinal cord injury. **A**. Representative images of 3 days post injury (3 dpi) at the epicenter of the injury. Spinal cord sections were immunostained for NG2 (green), GFAP (red) and counterstained with DAPI (blue). At the right are enlargements of the boxed areas in the images at the left. Scale bars = 100 μm. **B**. Quantified NG2 immunoreactivity at 3 dpi and, **C**. 7 dpi. At 3 and 7 dpi NG2 levels were significantly at the epicenter in the TG2-A-cKO mice. N=4 animals per group. ***p<0.001.

**Figure 4:**
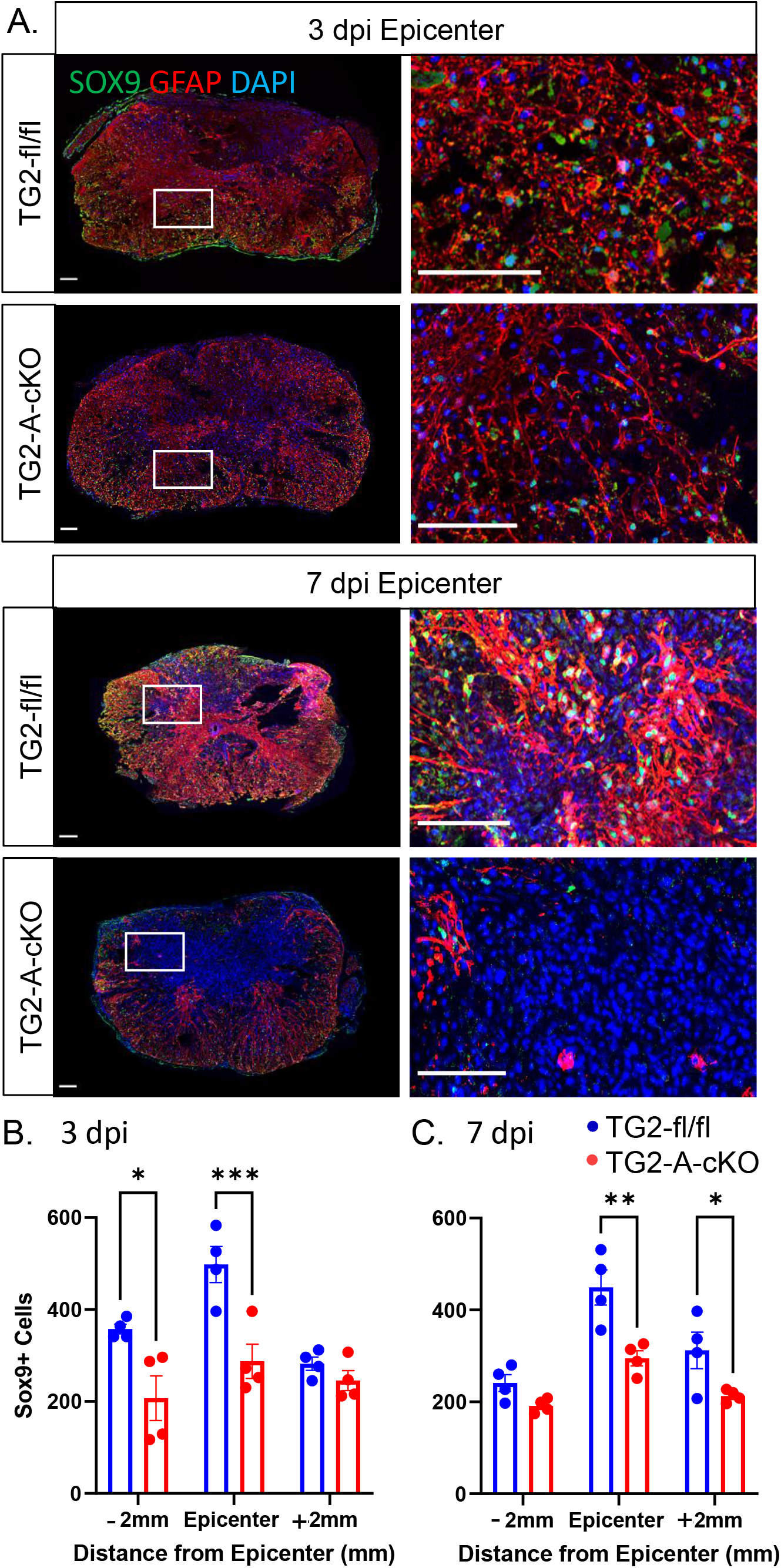
Deletion of TG2 from astrocytes results in a significant decrease in SOX9 positive cells after spinal cord injury. Spinal cord sections were immunostained for SOX9 (green), GFAP (red) and counterstained with DAPI (blue). **A**. Representative images of 3 and 7 days post injury (3 dpi, 7dpi) at the epicenter of the injury. Scale bars = 100 μm. The number of SOX9 positive cells was determined +/-2 mm from the site of injury as well as the epicenter at **B**. 3 dpi and **C**. 7 dpi. At 3 dpi the number of SOX9 positive cells in TG2-A-cKO mice was significantly decreased at both at -2mm and the epicenter. At 7 dpi there was still a significant decrease in the number of SOX9 cells at the epicenter in TG2-A-cKO mice, as well as at +2mm to the injury site. *p<0.05, **p<0.01, ***p<0.001. N=4 animals per group.

Lastly, to determine if there was increased neuron survival post SCI in the TG2-A-cKO mice, the number of NeuN positive cells were determined. At 3 dpi there was a slight but significant increase in the number of NeuN positive cells in the TG2-A-cKO mice caudal to the lesion center. However, by 7 dpi there was no significant differences in NeuN (**Fig. 5**).

**Figure 5:**
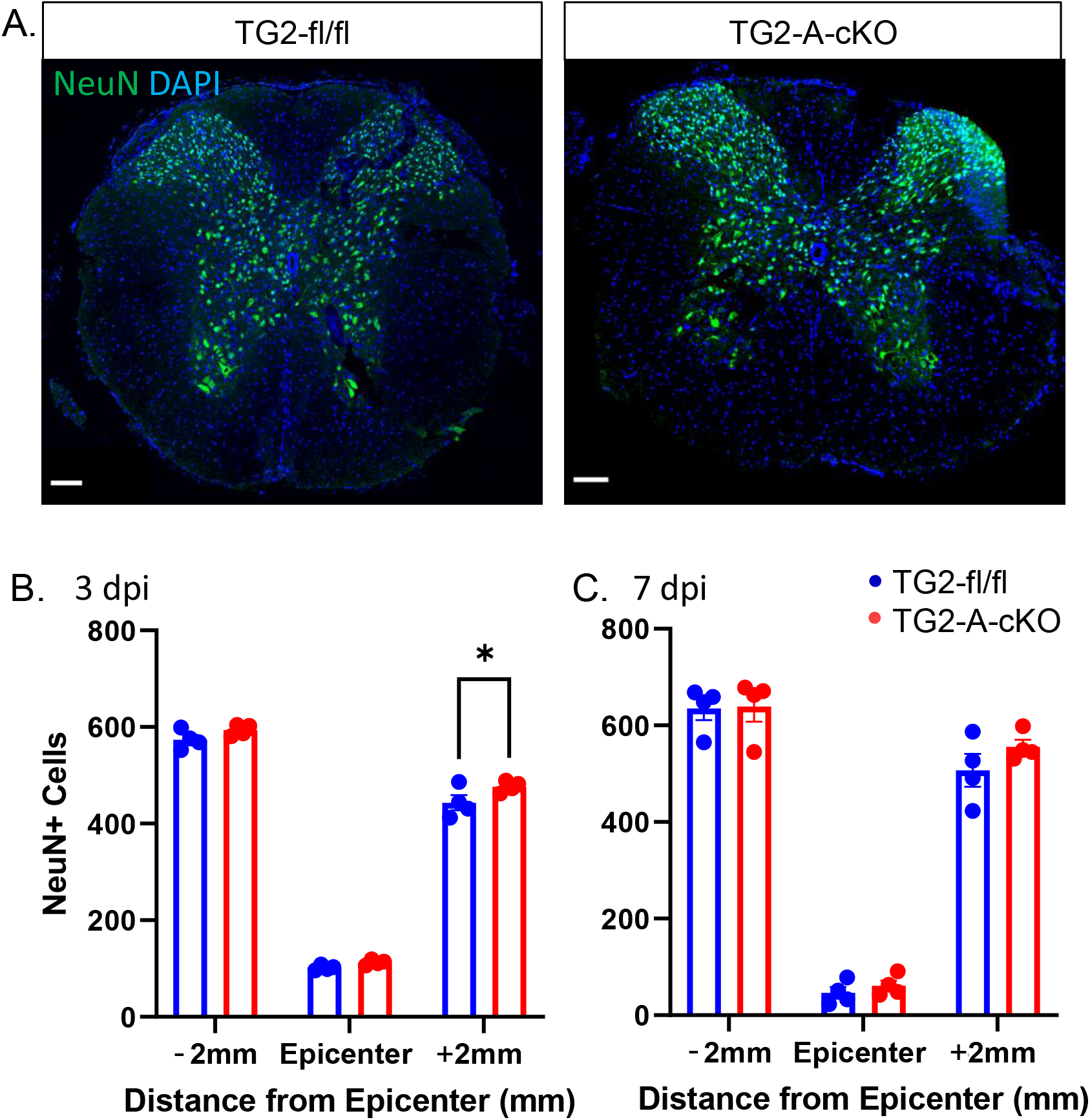
Deletion of TG2 from astrocytes did not result in a significant increase in neuron survival after spinal cord contusion injury. Spinal cord sections were immunostained for NeuN, a neuronal nuclear marker and the number of NeuN positive cells was determined +/-2mm from the site of injury as well as the epicenter. **A**. Representative images at 3 dpi 2 mm caudal to the epicenter of the injury. Scale bars = 100 μm. The number of NeuN positive cells was determined +/-2 mm from the site of injury as well as the epicenter at **B**. 3 dpi and **C**. 7 dpi. At 3 dpi the number of NeuN positive cells in TG2-A-cKO was significantly increased at +2mm (*p<0.05), however by 7 dpi there was no significant differences between the TG2-A-cKO and TG2fl/fl groups. N=4 animals per group.

### 3.3 Deletion of TG2 from astrocytes results in an upregulation of genes following SCI

To begin to understand how deletion of TG2 from astrocytes impacts gene expression after SCI, we carried out whole tissue RNA-seq. At 3 dpi RNA was extracted from 5 mm sections of the spinal cord centered on the site of injury from TG2-A-cKO and TG2fl/fl mice (n=4). Intriguingly, the 37 genes that were differentially expressed in the TG2-A-cKO mice after injury were all upregulated compared to TG2fl/fl mice (**Fig. 6**, Supplemental Table 1). Unexpectedly, gene ontology-based pathway analysis using Enrichr revealed 22 out of 39 GO terms of biological processes with an adjusted p-value of less than 0.05, are involved in lipid and fatty acid uptake, storage and metabolism (Supplemental Table 2). A STRING network representation shows the interconnectedness and preponderance of lipid metabolism pathways (**Fig. 7**). We then validated the expression of 3 key genes that are involved in lipid processing, lipoprotein lipase (Lpl), perilipin 1 (Plin1) and fatty acid binding protein 4 (Fabp4) via qPCR and found that all three were significantly upregulated in the spinal cords from the TG2-A-cKO mice 3 dpi (**Fig. 8**).

**Figure 6.**
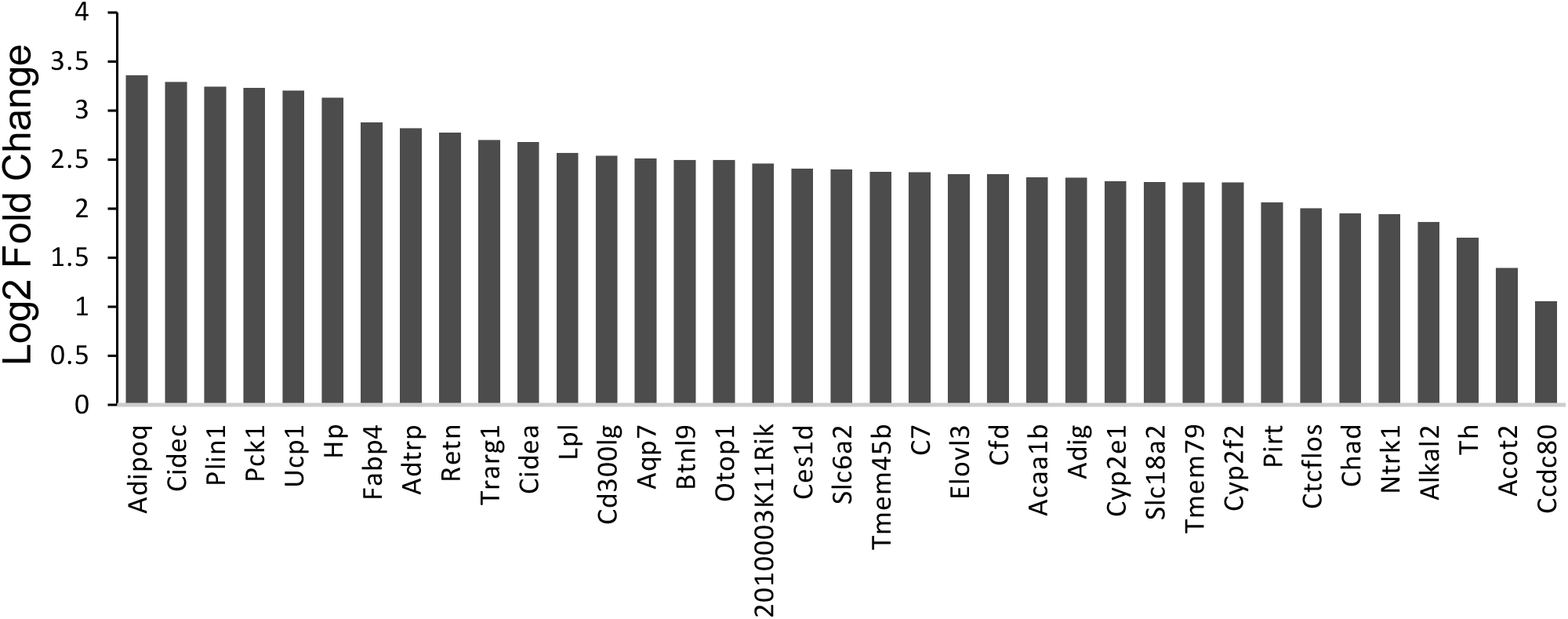
Deletion of TG2 from astrocytes results in an upregulation of genes involved in lipid uptake and metabolism following SCI. Five mm of spinal cord surrounding the lesions were collected from TG2-A-cKO and TG2fl/fl mice at 12–16 weeks of age (4 mice in each group) 3 days postinjury, RNA isolated and used for RNA-seq. Thirty-seven genes were increased in injured spinal cord from TG2-A-cKO compared to TG2fl/fl mice. Graphic representation of log fold changes in gene expression in spinal cords from TG2-A-cKO is shown.

**Figure7.**
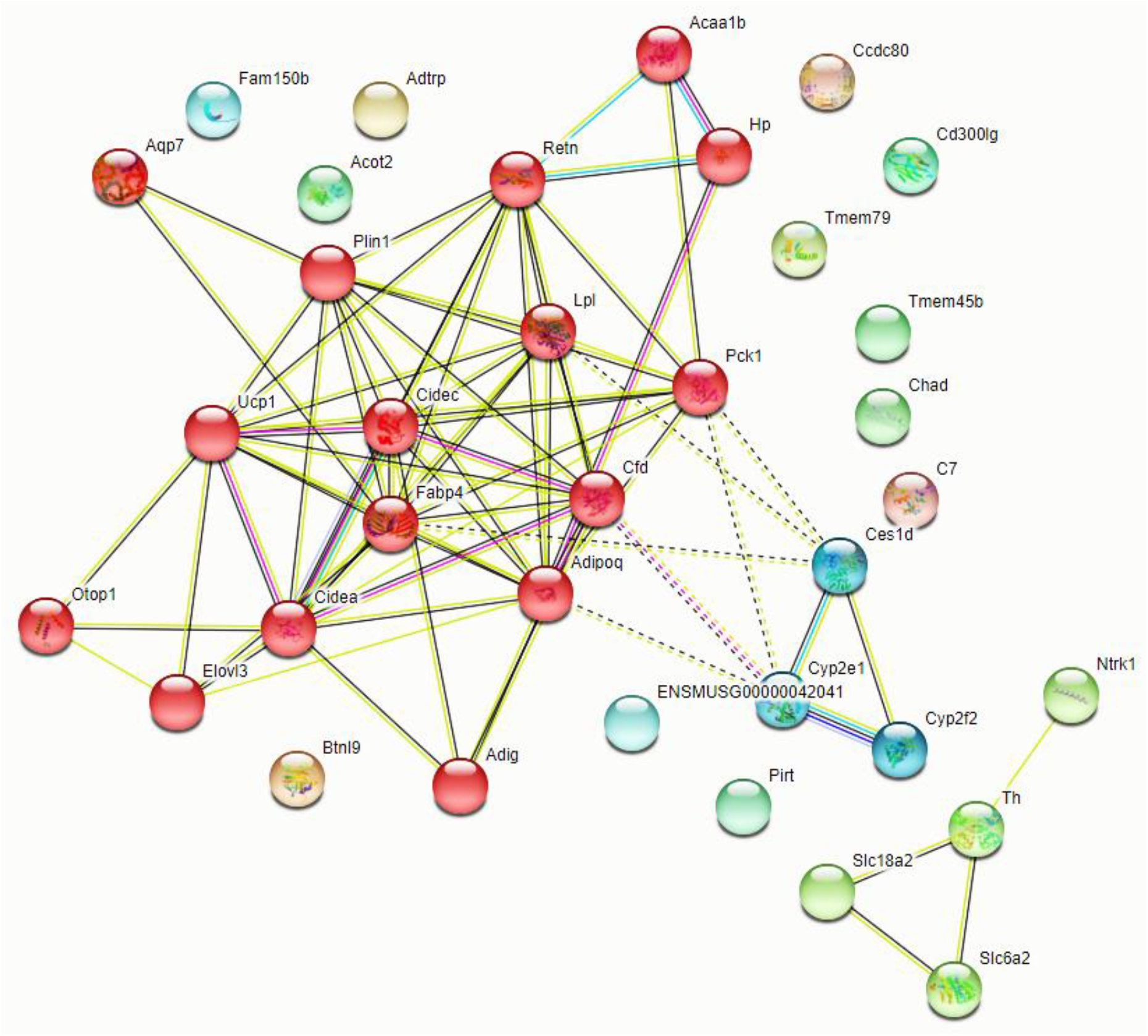
String analyses of genes that were upregulated in injured spinal cord from TG2-A-cKO compared to TG2fl/fl mice. Red hubs are genes that are directly involved in fatty acid metabolism, lipid storage and energy pathways.

**Figure 8:**
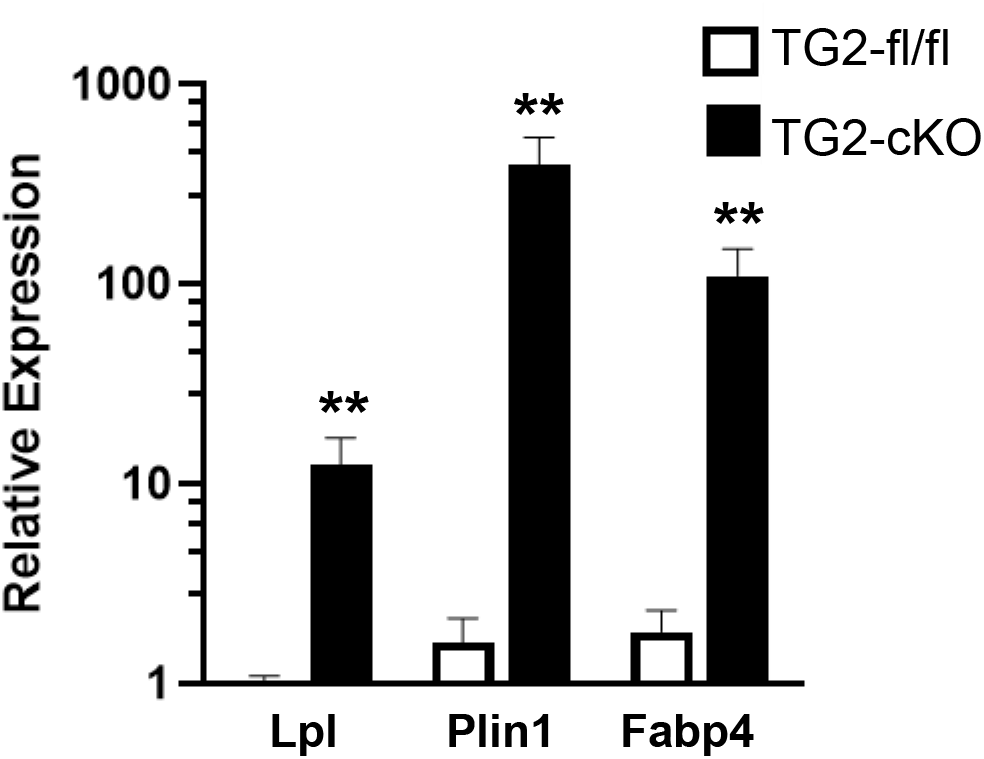
Deletion of TG2 from astrocytes increases the expression of genes that facilitate neuron-astrocyte metabolic coupling following SCI. TG2-A-cKO and TG2fl/fl mice at 12–16 weeks of age (4 mice in each group) were collected at 3 days postinjury, spinal cords removed and a 5 mm section of the spinal cord surrounding the lesion was used to prepare RNA followed by reverse transcription and for qPCR. Fatty acid binding protein 4 (Fabp4) is a carrier protein of fatty acids and likely facilitates the transport of fatty acids from neurons to astrocytes. Lipoprotein lipase (Lpl) degrades lipoprotein particles and thus upregulation would allow more fatty acids to enter the astrocytes and perilipin 1 (Plin1) increases the storage capacity of lipid droplets in astrocytes. Gapdh was used as a control. **p<0.01, unpaired t-test.

### 3.4 Treatment of wild type mice with the irreversible TG2 inhibitor VA4 significantly improves functional recovery after acute spinal cord injury

In a previous study we had found that treatment of astrocytes with VA4 phenocopied the effects of TG2 deletion, i.e., they were more resistant to OGD-induced cell death [13]. To determine if VA4 was also efficacious in vivo, we administered VA4 (or vehicle only as a control) to wild type mice starting at 1 hour post SCI and then 24 and 48 hours later. Remarkably, we found that by 7 dpi the VA4 treated mice showed significantly better motor function compared to mice treated with vehicle only (**Fig. 9**). As with our TG2-A-cKO mouse model, wild-type control animals do not recover past an average TMS score of 18-19, while VA4 treated animals achieve a recovery of 28 points, very similar to TG2-A-cKO animals (**Fig. 1**).

**Figure 9:**
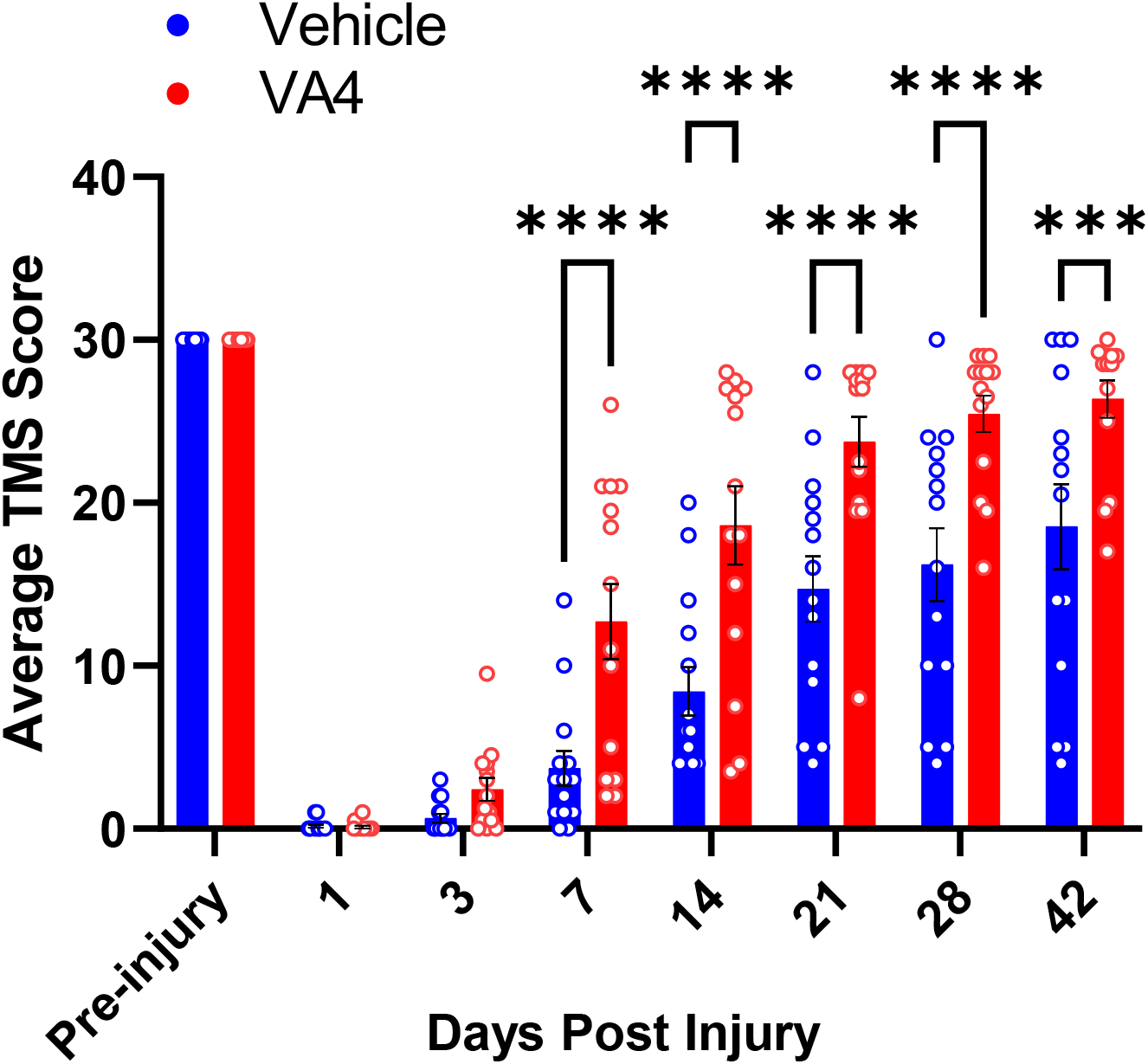
Treatment with the TG2 inhibitor VA4 improves functional recovery after spinal cord injury (SCI). Wild type mice at 12–16 weeks of age (14 mice in each group) were anesthetized and subjected to SCI as in Figure 1. Mice were administered VA4 at 15 mg/kg i.p. 1 hour post injury and then at 24 and 48 hours. Control animals received Cremaphor only. Motor function was assessed prior to injury and at the days indicated post-injury using the Toyama mouse score (TMS) to assess locomotor function [25]. Data are expressed as an average of the right hind limb and left hind limb. Data were analyzed by two-way ANOVA with Bonferroni’s multiple comparisons test, ***p<0.001, ****p<0.0001. TG2-fl/fl

## 4. Discussion

Astrocytes play an indispensable role in determining outcomes subsequent to SCI. When the CNS is injured, astrocytes undergo a process called reactive astrogliosis [4]. Acutely, this response is a defensive reaction aimed at limiting tissue damage. However, astrogliosis often results in processes that may inhibit functional recovery or fail to provide adequate support to promote axonal outgrowth [4, 5, 32-34]. Conversely, transplantation of non-reactive astrocytes can promote recovery after SCI [35-37] and in an animal model of Parkinson neurodegeneration [38]. In the injured CNS, astroglial response is a highly regulated process that is characterized by changes in astrocyte gene expression, morphology, proliferative state, and mobility, and is dependent upon numerous factors such as location, severity and time after insult [1]. Astrocytic modulatory proteins play key roles in determining how they respond to injury and the extent to which they inhibit or support functional recovery. Identifying and understanding the role different modulatory proteins play in the injury response is essential for the development of intervention strategies to promote the ability of reactive astrocytes to support and not hinder functional recovery. One such modulatory protein that plays an important role in regulating astrocyte function and response to injury is TG2.

Previously we demonstrated that deletion of TG2 from astrocytes or treatment with the selective TG2 inhibitor VA4, significantly increases their survival in OGD conditions [6, 13]. Deletion of TG2 from astrocytes also significantly increases their ability to protect neurons from ischemic-induced death [6, 39]. Given the fact that TG2 regulates transcriptional processes [40, 41], we carried out RNA-seq on wild type and TG2-/-astrocytes and found that pro-survival/pro-regenerative genes were upregulated in the TG2-/-astrocytes [39]. Although these findings indicate that deletion of TG2 from astrocytes is beneficial in the context of CNS injury, this needs to be tested in an in vivo model. Therefore, we used TG2-A-cKO and TG2fl/fl mice [39] in a contusion SCI model followed by analysis of motor function over a 42 day period. Remarkably, motor function in the mice with astrocytic TG2 deleted was significantly better starting at 7 dpi and continuing through 42 dpi. In fact, by 35 dpi many of the TG2-A-cKO mice had recovered most of their motor function based on the TMS score (**Fig. 1**). These data clearly indicate that in vivo deletion of TG2 from astrocytes enhances their supportive role post traumatic injury.

Immunohistochemical analyses of the spinal cords from the TG2-A-cKO and TG2fl/fl mice at 3 and 7 dpi suggests a modulation of the glial scar in animals with astrocytic deletion of TG2. In agreement with what we had observed previously [39], GFAP immunoreactivity as a general indicator of astrogliosis was significantly reduced in the TG2-A-cKO mice. Likewise, we find a significant reduction in the expression of the neurite growth inhibitory CSPG NG2 and the upstream regulator of inhibitory CSPGs, Sox9. Sox9 is associated with negative outcomes following CNS injury as it upregulates inhibitory CSPGs as well as other genes associated with glial scar formation [15], including NG2 (Cspg4) [31]. Deletion of Sox9 reduces inhibitory CSPG levels, increases reactive sprouting and improves outcomes after a contusion SCI [16, 17]. In line with these previous studies, and the functional recovery observed in TG2-A-cKO mice, the number of Sox9 positive cells and NG2 immunofluorescence were significantly reduced in the TG2-A-cKO mice at both 3 and 7 dpi compared to TG2fl/fl mice. It has been suggested that NG2 plays a role in preventing axon regeneration across newly formed glia scars [34]. An immunohistochemical analyses of post mortem samples from control and injured/lesioned human spinal cords revealed that NG2 was present in the evolving astroglial scar and therefore it was suggested that it played a role in inhibiting successful CNS regeneration [14]. While the direct role of reactive astrocytes in the production of CSPGs remains unclear [32], our finding of reduced Sox9 and NG2 strongly indicate that deletion of TG2 from astrocytes results in an environment that is more permissive for successful regeneration following a CNS injury.

To begin to understand how deletion of TG2 from astrocytes impacts the genetic landscape following SCI thereby promoting regeneration, we carried out RNA-seq on spinal cords from TG2-A-cKO and TG2fl/fl mice 3 dpi. Thirty-seven genes were differentially expressed in the TG2-A-cKO spinal cords following SCI and all were upregulated. This is in line with the finding that when TG2 is deleted from astrocytes the majority of genes that are differentially expressed are upregulated [39]. Using the FunRich interaction network analysis tool [27] (See Supplemental Figure S2), it was found that the upregulated genes were primarily involved in energy pathways, fatty acid metabolism and lipid storage, which is also illustrated in the STRING analysis (**Fig. 7**) and gene ontology analysis with Enrichr (Supplemental Table S2). At first glance these results were puzzling; however, upon further investigation these results are quite exciting given that astrocyte-neuron metabolic coupling is crucial for maintaining healthy neurons. SCI induces neuronal hyperactivity along with glial activation creating a maladaptive environment [42] as hyperactive neurons release toxic fatty acids. These fatty acids need to be transferred to astrocytes in lipid particles for astrocytes to protect the neurons [19] and both Plin1 and Lpl, which are significantly increased in injured TG2-A-cKO spinal cords, would increase fatty acid flux into astrocytes. Lpl plays a key role in the degradation of triglyceride particles and release of fatty acids which plays a significant role in maintaining energy balance in astrocytes and neurons [18]. At the same time, increased Plin1 expression would increase the storage capacity of lipid droplets in astrocytes [43] which would protect neurons from oxidative stress, and promote a more regenerative environment [44]. Further, prior studies have shown that the ability to transport lipids to astrocytes for lipid droplet formation is essential to maintain healthy neurons [20]. Overall, it is clear that astrocyte-neuron metabolic coupling is likely a critical factor in the regenerative processes that take place after SCI and that upregulation of genes that mediating the coupling in the TG2-A-cKO may contribute to the improve functional outcomes from SCI. Therefore, TG2 inhibition/ablation in astrocytes does not prevent reactive gliosis, but rather promotes a more regenerative phenotype.

In vitro both deletion of TG2 from astrocytes and treatment with VA4 significantly attenuated OGD-induced cell death [13]. Therefore, the final aspect of this study was to determine if the TG2 inhibitor VA4 would improve motor function after SCI in a manner similar to what was observed with astrocytic TG2 deletion. For these studies wild type mice were administered either 15 mg/kg VA4 or vehicle 1 hour after SCI and then again at 24 and 48 hours. Remarkably, this simple 3 dose paradigm with VA4 significantly improved motor function. VA4 is an irreversible inhibitor that bears an acrylamide “warhead” which is quickly becoming the most popular warhead [45] in novel targeted covalent inhibitors on the market.. What is critically important to the specificity of the drug is the “targeting domain”, which in the case of VA4 is highly specific for TG2; VA4 does not inhibit any other transglutaminase tested [45]. With a low toxicity profile towards neurons and high potency for inhibiting the transamidating activity of TG2 [21], VA4 has therapeutic potential. These findings suggest that selective, irreversible TG2 inhibitors have the potential to be considered for the treatment of spinal cord and other acute CNS injuries. Future studies will continue to explore the mechanisms of action and window of treatment for the use of VA4 in improving outcomes following SCI.

## Supplementary Materials

Figure S1: Supplementary code for fluorescence quantification. Figure S2: Functional analysis of genes that are significantly upregulated in TG2-A-cKO mice 3 days postinjury using FunRich (http://www.funrich.org). Table S1: Genes that are significantly different in spinal cords of GFAP-Cre+/-TG2fl/fl mice compared to TG2fl/fl mice 3 days postinjury. Table S2: Functional analysis of genes that are significantly upregulated in TG2-A-cKO mice 3 days postinjury using Enrichr.

## Author Contributions

Conceptualization, GVWJ, CP, AE, JR; Experimentation:, GVWJ, CP, AE,JR, GS, VV, JE, PG; VA4 synthesis: JWK; Data Analysis, AE, JE, PG, AV, GS, CP.; Manuscript preparation, Review & Editing, GVWJ, CP, AE, JWK; Funding Acquisition, GVWJ, CP.

## Institutional Review Board Statement

The study was conducted according to the guidelines of the Declaration of Helsinki, and approved by the Institutional Review Board of University of Rochester (UCAR) (protocol #s: APP 2007-023E [5/14/2021] and APP 2010-07E [9/16/2019]).

## Acknowledgements

RNA-seq was performed on a fee for service basis by the University of Rochester Genomics Research Center (GRC.) The authors would like to thank Dr. Michael Welte for his insightful comments about the RNA-seq analyses.

## Figure Legends

**Supplemental Figure S1.**
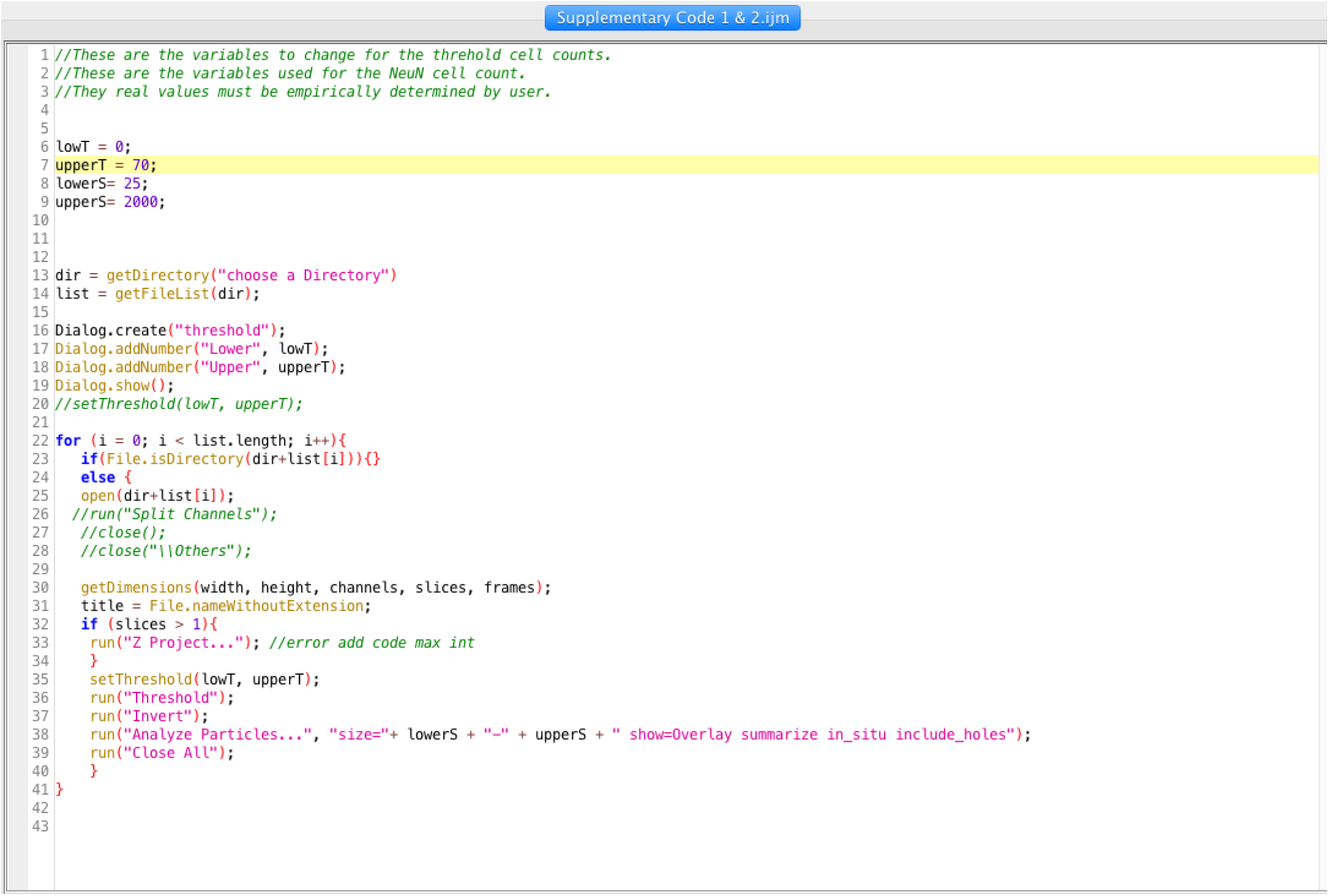
Supplemental code for Fiji Image analysis used for immunofluorescence quantification.

**Supplemental Figure S2.**
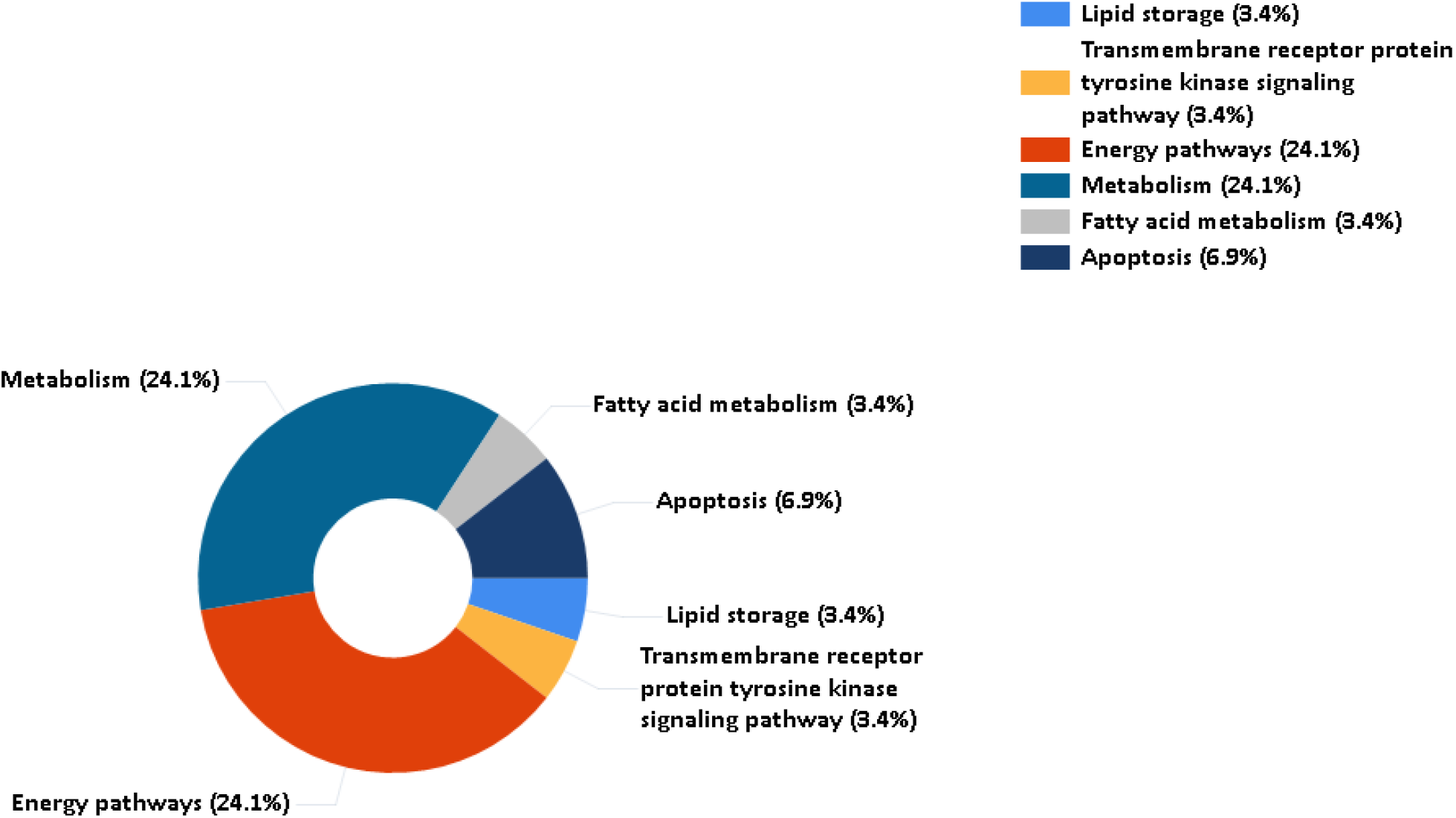
Functional analysis of genes that are significantly upregulated in GFAP-Cre+/-TG2fl/fl mice three days post injury using FunRich (http://www.funrich.org).

**Supplemental Table 1:**
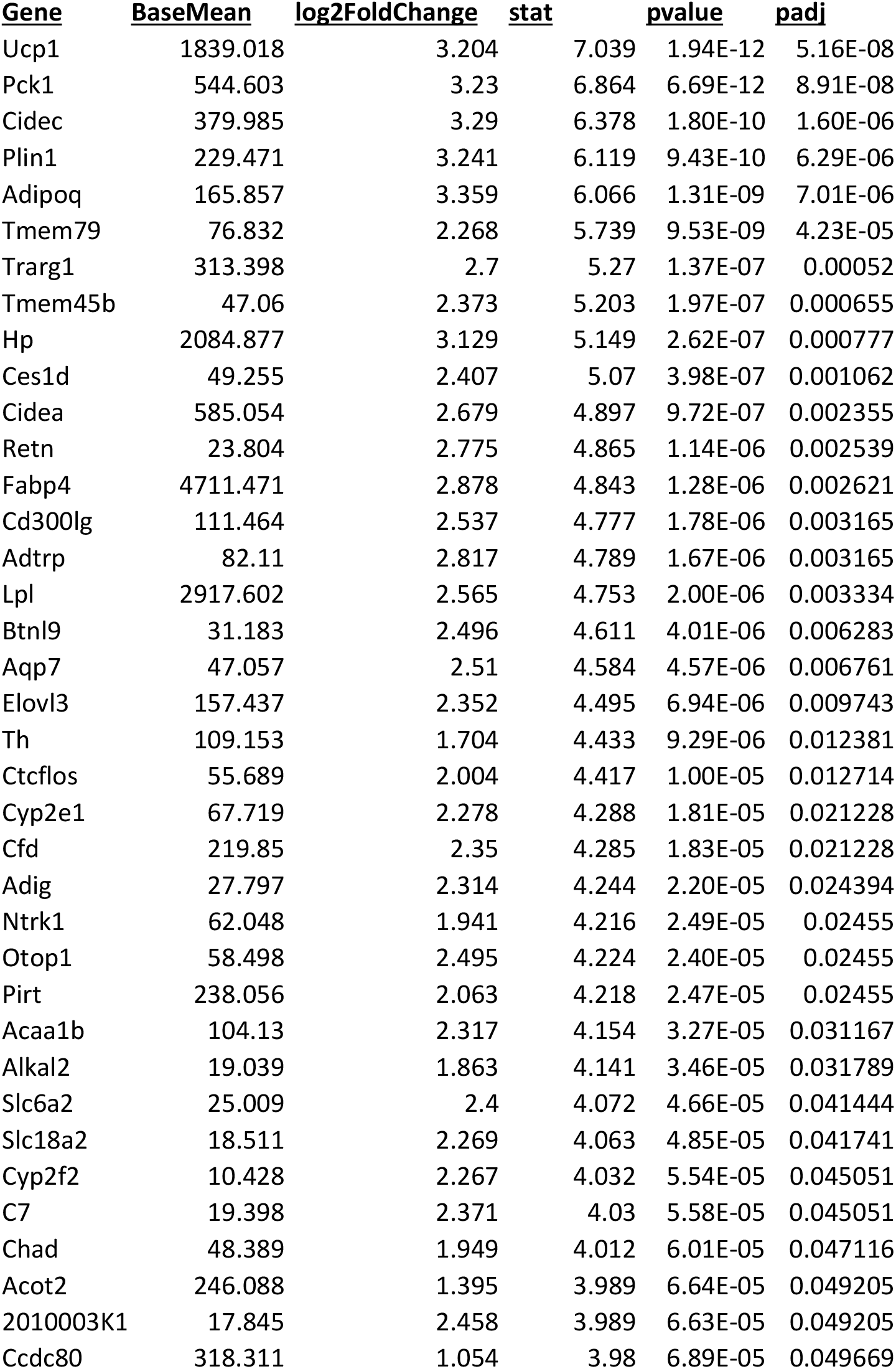
Genes that are signficantly different in spinal cords of GFAP-Cre+/-TG2fl/fl mice compared to TG2fl/fl mice 3 days postinjury.

**Supplemental Table 2.**
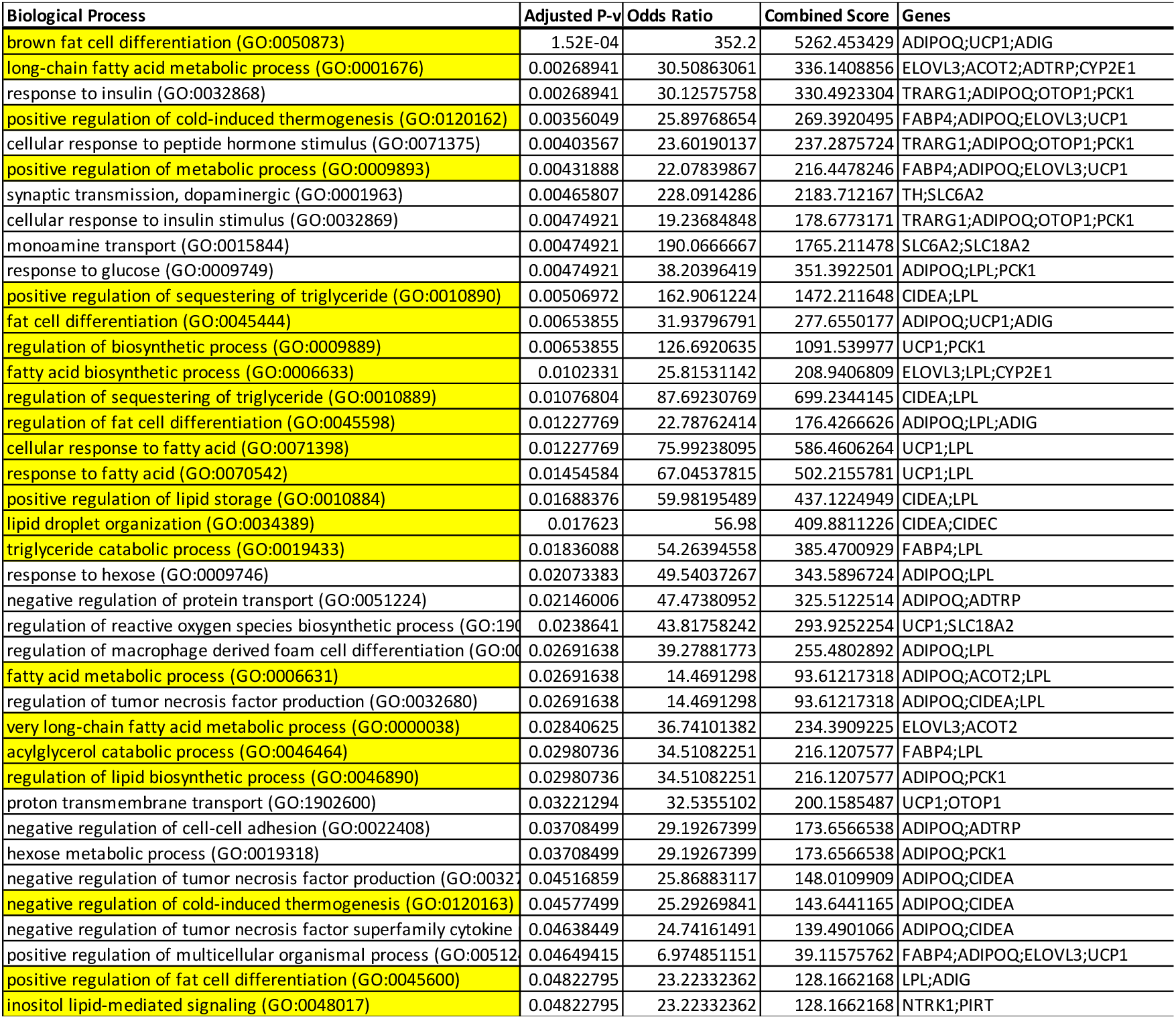
Gene ontology pathway analysis identifies a preponderance of lipid biosynthetic pathways among differentially expressed genes in mice lacking astrocytic TG2. 22 out of 39 GO terms with an adjusted p-value of less than 0.05 are involved with lipid biosynthetic pathways and storage (highlighted in yellow). Enrichr, GO Biological Process 2021.

